# Closed mitosis requires local disassembly of the nuclear envelope

**DOI:** 10.1101/779769

**Authors:** Gautam Dey, Siân Culley, Scott Curran, Ricardo Henriques, Wanda Kukulski, Buzz Baum

## Abstract

At the end of mitosis, eukaryotic cells must segregate both copies of their replicated genome into two new nuclear compartments (1). They do this either by first dismantling and later reassembling the nuclear envelope in a so called “open mitosis”, or by reshaping an intact nucleus and then dividing into two in a “closed mitosis” (2, 3). However, while mitosis has been studied in a wide variety of eukaryotes for over a century (4), it is not known how the double membrane of the nuclear envelope is split into two at the end of a closed mitosis without compromising the impermeability of the nuclear compartment (5). In studying this problem in the fission yeast *Schizosaccharomyces pombe*, a classical model for closed mitosis (5), we use genetics, live cell imaging and electron tomography to show that nuclear fission is achieved via local disassembly of the nuclear envelope (NE) within the narrow bridge that links segregating daughter nuclei. In doing so, we identify a novel inner NE-localised protein Les1 that restricts the process of local NE breakdown (local NEB) to the bridge midzone and prevents the leakage of material from daughter nuclei. The mechanics of local NEB in a closed mitosis closely mirror those of NEB in open mitosis (3), revealing an unexpectedly deep conservation of nuclear remodelling mechanisms across diverse eukaryotes.

## Introduction

A key event in the process of cell division in eukaryotes is the partitioning of the nuclear genome into two nuclear compartments. To achieve this, replicated sister chromosomes detach from the inner nuclear envelope (INE) (3), enabling them to be separated from one another through the work of a microtubule-based spindle (6), before being sorted into two new, physically separate nuclei at mitotic exit. Eukaryotic cells have adopted a wide spectrum of strategies to coordinate nuclear remodelling with chromosome segregation (2). At one extreme, in a so called “open mitosis”, cells first disassemble the nuclear lamina and the continuous nuclear envelope (NE) at mitotic entry, and then reverse this process by reassembling the structure around separated chromosomes at mitotic exit. At the other extreme, in a “closed mitosis”, because the nuclear/cytoplasmic compartment barrier remains intact, spindle poles must be inserted into the nuclear envelope (7) to form an intranuclear spindle that can drive chromosome segregation. This spindle is then disassembled as the NE is divided into two (8). While these different modes of nuclear division share key features, and despite there being a range of intermediate states (9, 10), the resolution of a closed mitosis presents a unique topological challenge. Currently it is not understood how this is overcome to enable organisms undergoing a closed mitosis to divide the double nuclear envelope without compromising nuclear integrity.

To shed light on this process, we chose to study nuclear division in the fission yeast, *Schizosaccharomyces pombe* (*S. pombe*), which serves as an experimentally tractable example of an organism that undergoes a classic closed mitosis. Previous studies have shown that the *S. pombe* nucleus does not tear at mitotic exit (11), as it does in the related yeast *S. japonicus*. Instead the nucleus constricts to form a dumbbell shape with a thin nuclear bridge around the anaphase spindle (5). While the organisation and dynamics of the anaphase spindle have been studied in some detail (12–14), it is not known how the nuclear envelope is then remodelled to generate two new nuclei at the end of this process without compromising the nuclear-cytoplasmic compartment barrier.

## Results

To explore this question, we used a synthetic nuclear-localised GFP construct to characterise the dynamics of nuclear fission, and to determine the extent to which the nuclear permeability barrier is maintained throughout the process (Figure 1a, 1b). Although nuclear GFP levels remained constant throughout the division process (as expected for a closed mitosis), we observed a gradual loss of GFP from the nuclear bridge prior to nuclear division (Figure 1b, using an automated analysis pipeline described in Figure S1b and Methods). Importantly, this occurred without the visible leakage of GFP from daughter nuclei (Figure 1a). We then searched for a marker to examine nuclear envelope remodelling during the same time period. Doing so is complicated by the fact that in fission yeast, as in other eukaryotes, the outer face of the nuclear envelope is linked to the endoplasmic reticulum (ER) (15). In *S. pombe*, these take the form of thin ER tubules that bridge the outer NE and the ER sheets that underlie the plasma membrane (Figure 1c and Figure S1a). Since the ER is actively remodeled at the cell equator during mitosis (Figure S1a), it is difficult to differentiate nuclear envelope dynamics from bulk ER dynamics. However, in screening for a suitable marker we identified a hitherto-uncharacterised LEA domain (16) protein, SPAC23C4.05c, that localises exclusively to the nucleoplasmic surface of the inner NE throughout the cell cycle (Figure 1c, Figure S1c) without marking ER tubules or the cortex. Strikingly, SPAC23C4.05c was also seen concentrated at the stalk of each daughter nucleus during anaphase - a phenotype for which we named the protein Les1, or LEA domain protein Enriched in Stalks.

**Fig. 1.**
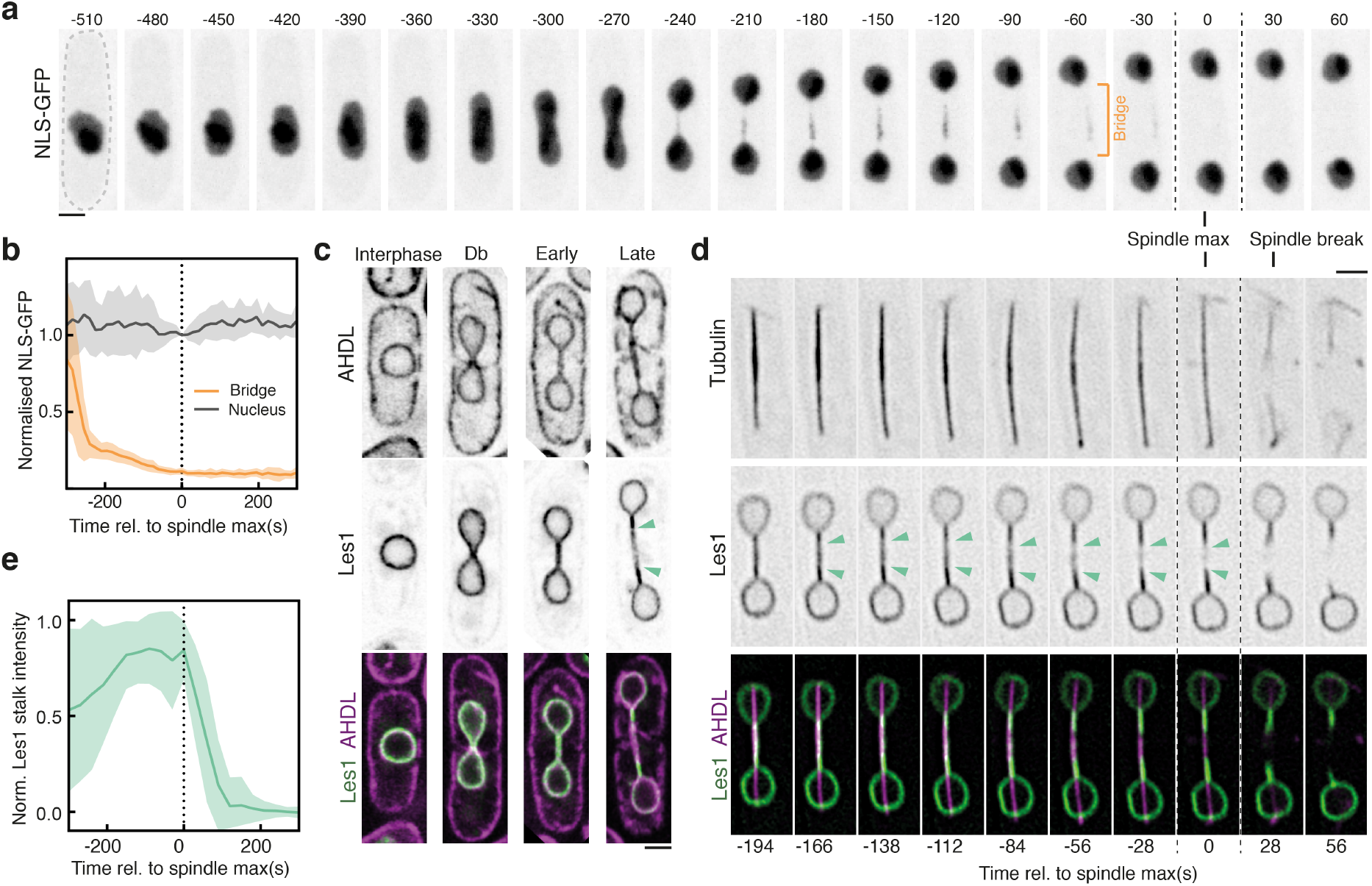
Les1 domains define the site of nuclear division. a. *S. pombe* cell (dotted outline) expressing a synthetic NLS-GFP construct undergoing mitosis; panels are maximum intensity projections of confocal images acquired every 30 seconds. Dotted vertical lines represent maximum spindle length (Spindle max, t=0). b. Mean (darker lines) and standard deviations (lighter bands) of averaged single-cell traces (between 11 at t=300 and 19 at t=0, from 2 strains) aligned at spindle max (t=0) and normalised to daughter nuclear intensity at t=0. c. Single Airyscan reconstructions of live cells at various stages of the cell cycle expressing a synthetic ER-localised mCherry construct (AHDL) and Les1 (Les1; SPAC23C4.05c) tagged with mNeonGreen at the endogenous locus. Green arrows mark the boundaries of the Les1 stalks, visible in late anaphase. d. Reconstructions using Super-Resolution Radial Fluctuations (SRRF) at 28 second intervals on single confocal slices, of a cell undergoing anaphase, expressing Atb2-mCherry (Tubulin) and Les1-mNeonGreen tagged at the endogenous loci. Green arrows mark Les1 stalk boundaries, which first become visible in mid-anaphase. e. Les1 intensity in stalks (1.5 µm from nuclear periphery) over time: mean (darker line) and standard deviations (lighter band) of between 11 (t=300) and 36 (t=0) single-cell traces, aligned at spindle max (t=0) and normalised to maximum bridge intensity. All scale bars = 2 µm.

Using Les1 as a marker, we were then able to follow the dynamic changes in nuclear shape that accompany spindle elongation in detail – as a single nucleus divides into two via a characteristic dumbbell-shaped intermediate (Figure 1d). The kinetics of spindle elongation are highly reproducible ((17), allowing us to align single-cell trajectories to the time point at which the spindle reaches its maximum length (see also Methods). At early stages of bridge formation, Les1 was found to concentrate in stalks (Figure 1d-e). At maximum spindle elongation, Les1 was visibly depleted from the midzone of the bridge (Figure 1d); a process that was followed, within a few seconds, by the breakage of the spindle (Figure 1d and Figure S1d).

Since these observations pointed to the midzone of the bridge as the site where nuclear fission occurs, we used correlative light microscopy and electron tomography of Les1mNeonGreen/mCherry-Atb2 dual-labeled cells (Figure 2a) to characterise early and late nuclear bridges (Fig 2b, 2c and Fig S2a). In early bridges, the nuclear envelope was seen bounding the spindle (narrowing at the base of the stalks and widening towards the midzone) and was studded with nuclear pores (Figure 2b). At an intermediate stage, the nuclear pores were completely excluded from the stalk and clustered in a central bulge (Figure S2b). By contrast, in late stage bridges, while the nuclear envelope still enveloped the spindle within stalks, there was no evidence of a continuous nuclear envelope within the midzone region of the bridge. Instead, spindle microtubules were seen projecting out of the two newly formed daughter nuclei, through stalk membranes that lacked nuclear pores (Figure 2b), into the cytoplasm (Figure 2c). All that was left of the central part of the nuclear bridge at this stage were fragments of ER membrane - an observation that explains the loss of nuclear GFP from the midzone of the late anaphase bridge (Figure 1).

**Fig. 2.**
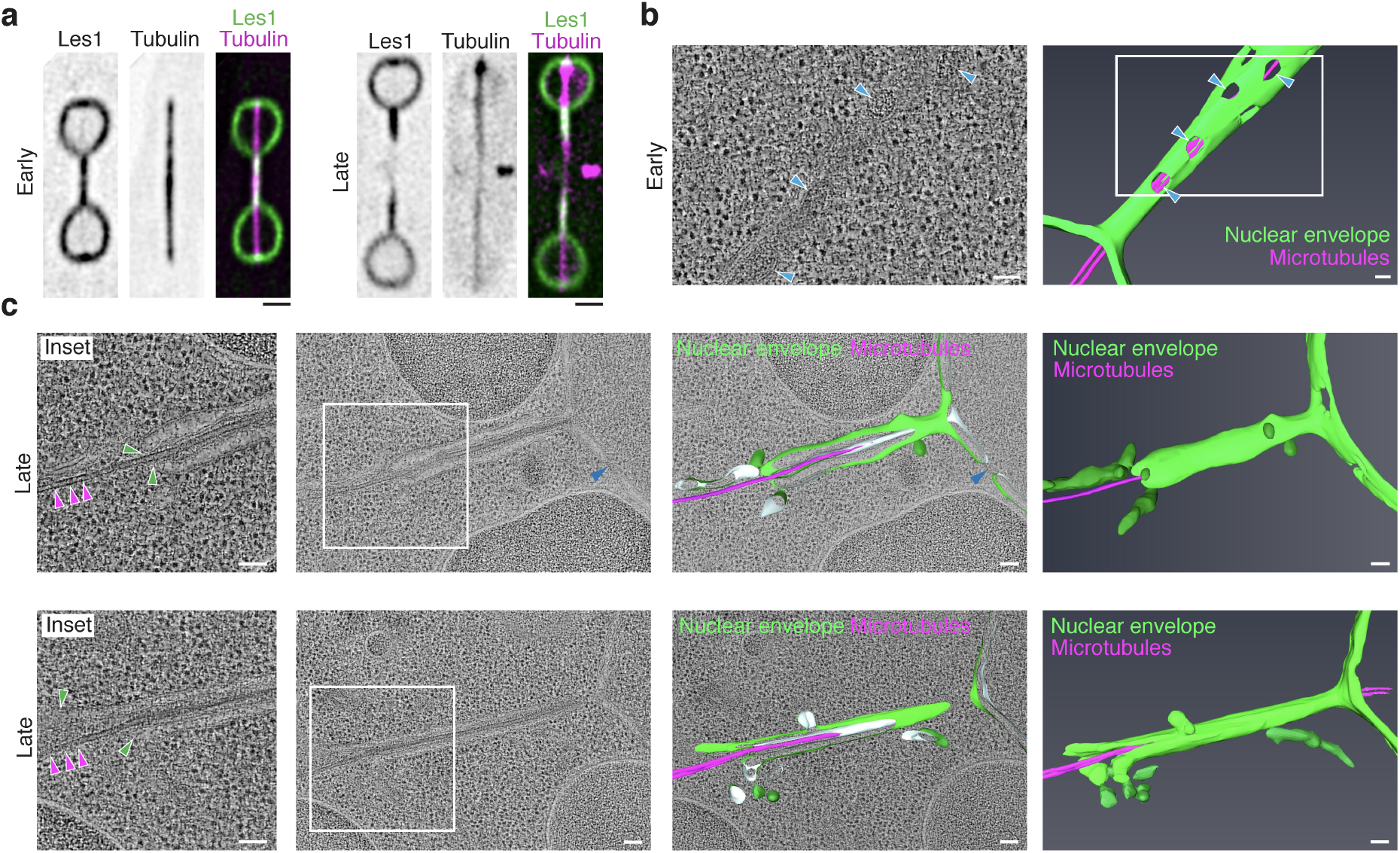
Nuclear division occurs by local nuclear envelope breakdown. a. Representative SRRF reconstructions from single confocal slices of cells in early (left-hand panels) and late (right-hand panels) anaphase, expressing Les1-mNeonGreen and Atb2-mCherry (tubulin) tagged at the endogenous loci. Scale bar = 2 µm. Note that intact spindle microtubules still persist in the late anaphase bridge despite midzone clearance of Les1. b. Virtual slice through electron tomogram (left) and 3D segmentation model (right) of early anaphase bridge (nuclear envelope in green, microtubules in magenta). White region indicates magnified region in left panel. Blue arrows indicate nuclear pores. Scale bars = 100 nm. c. Clipped and full view of 3D segmentation models (right panels) of late anaphase bridges (nuclear envelope in green, microtubules in magenta), superimposed on virtual slices through electron tomograms (second from left), magnified region indicated by white inset. Magenta arrows indicate microtubules, green arrows indicate stalk tip and limit of intact nuclear envelope. Blue arrows indicated nuclear pores. Scale bars = 100 nm.

If the nuclear envelope in the central region of the bridge is disassembled to induce nuclear division, as suggested by this unexpected observation, how is this midzone region itself specified? A clue to this came from the observation made using EM that nuclear pores are absent from late-stage bridges (Figure 2c).

In line with this, when we used light microscopy to track transmembrane (Cut11; hereafter “transmembrane”) and nucleoplasmic ring (Nup60; hereafter “peripheral”) components of the nuclear pore complex (NPC) through mitosis (nucleoporins or “Nups”), we observed NPCs being depleted from stalk regions of the bridge where Les1 accumulated (Figure 3a). At the same time, the excluded NPCs became concentrated within the middle region of the bridge, as previously reported (18, 19), where Les1 levels appeared depleted (Figure 3a and 3b). Thus, Les1 and NPCs appear to have a mutually exclusive localisation. In line with this, occasional ‘stray’ NPC clusters located distal to the midzone correspond to areas of local Les1 depletion within the stalk region (Figure 3c, 3d) and local bridge dilation (Figure S3a).

**Fig. 3.**
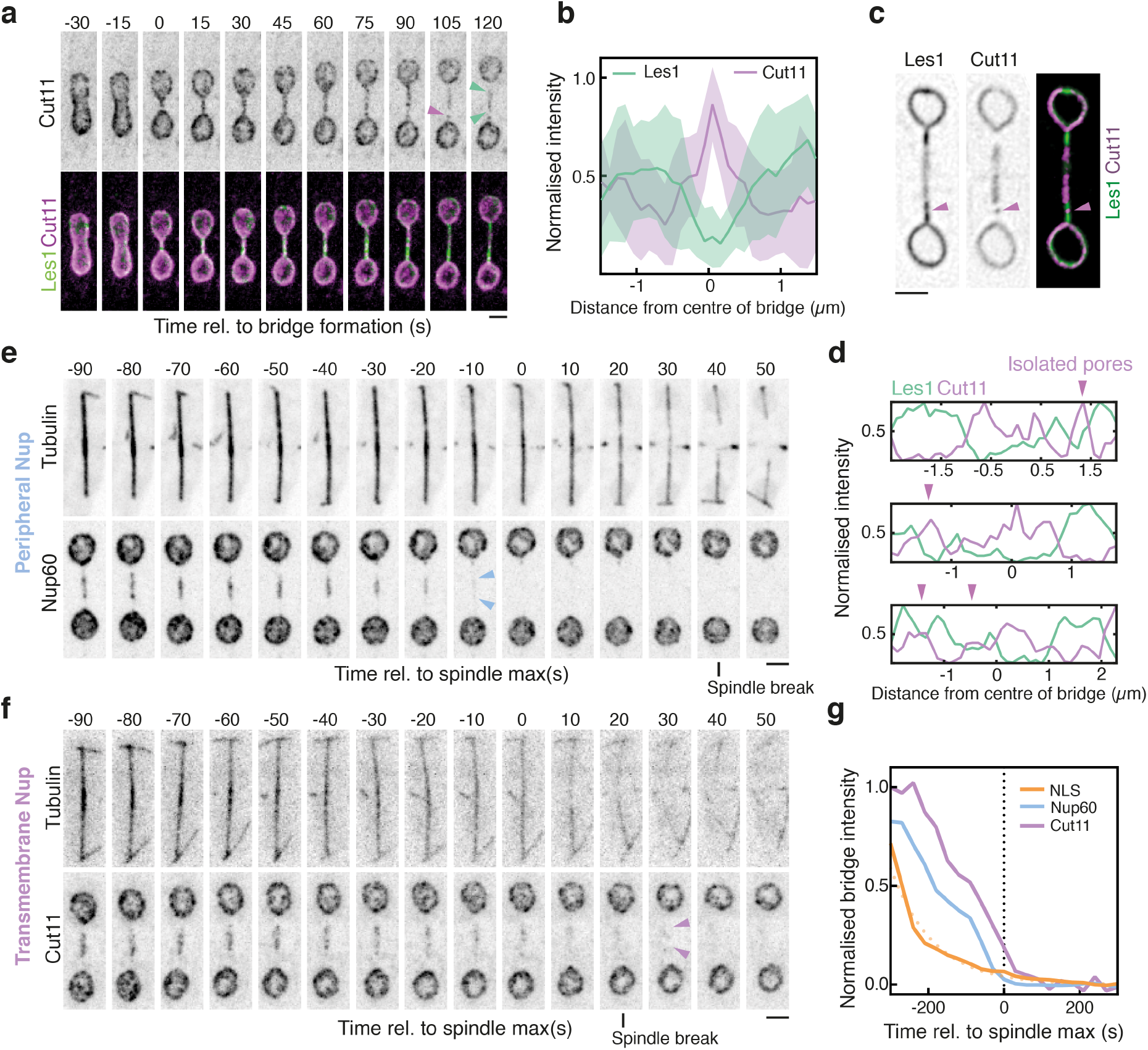
Les1-restricted stepwise NPC disassembly drives local NEB. a. Maximum intensity projections of spinning disk confocal images acquired every 15 seconds of S. pombe expressing Les1-mNeonGreen and Cut11-mCherry tagged at the endogenous loci. Bridge formation is at t=0. Green arrows mark boundaries of Les1 stalks, magenta arrow indicates stray pore cluster. Scale bar = 2 µm. b. Averaged line traces (darker lines = mean, lighter bands = standard deviation) of Les1 and Cut11 intensities along the bridge for 16 cells at bridge length 3 µm. c. SRRF reconstruction from single confocal images of a cell expressing Les1-mNeonGreen and Cut11-mCherry. Magenta arrow indicates stray nuclear pore cluster, with 4 more examples marked upon the line scans along the bridges of 3 illustrative cells in d. Maximum intensity projections of confocal images of cells expressing either Nup60-mNeonGreen (e) or Cut11-GFP (f) along with Atb2-mCherry and acquired at 10 second intervals. Scale bars = 2 µm. g. Averaged normalised intensity traces for NLS-GFP (between 22 cells at t=-300 to 37 at t=0), Nup60-mNeonGreen (between 11 cells at t=-300 to 34 at t=0), and Cut11-mCherry (between 11 cells at t=-300 to 34 at t=0), aligned by spindle max (t=0). Dotted yellow line indicates single exponential fit to NLS-GFP average. Data from two strains were combined based on the analysis described in Figure S3b

In an open mitosis, the stepwise removal of NPCs leads to fenestration and loss of structural integrity during local nuclear envelope breakdown (10). We observed precisely this sequence of events in *S. pombe*: the NPC signal in the bridge was gradually lost after dumbbell collapse (Figure 3e-f), with membrane Nups being lost last, as the nuclear NLS-GFP signal was seen disappearing from the midzone (Figure 3g). Within a single bridge, fast imaging revealed that distinct clusters of NPCs disappeared at different times, though with the relative sequence preserved (Figure S3d). Importantly, the complete loss of membrane Nups occurred prior to, and was independent of, cell division itself (Figure S3e-f).

These data strongly suggest that NPCs, and the subsequent phenomenon of nuclear envelope fragmentation, are restricted to the bridge midzone by Les1. To test this idea, we examined the phenotype of *les1*Δ cells. Strikingly, the spatial organization of NPCs in the bridge was completely lost in this deletion strain. NPC components were found distributed uniformly both in early and late stage bridges of *les1*Δ. cells (Figure 4a). This was associated with a change in the position at which the bridge broke. Thus, the spindle broke in a Les1-depeleted zone at the centre of the wildtype bridge, but at random locations in *les1*Δ cells (Figure 4b).

**Fig. 4.**
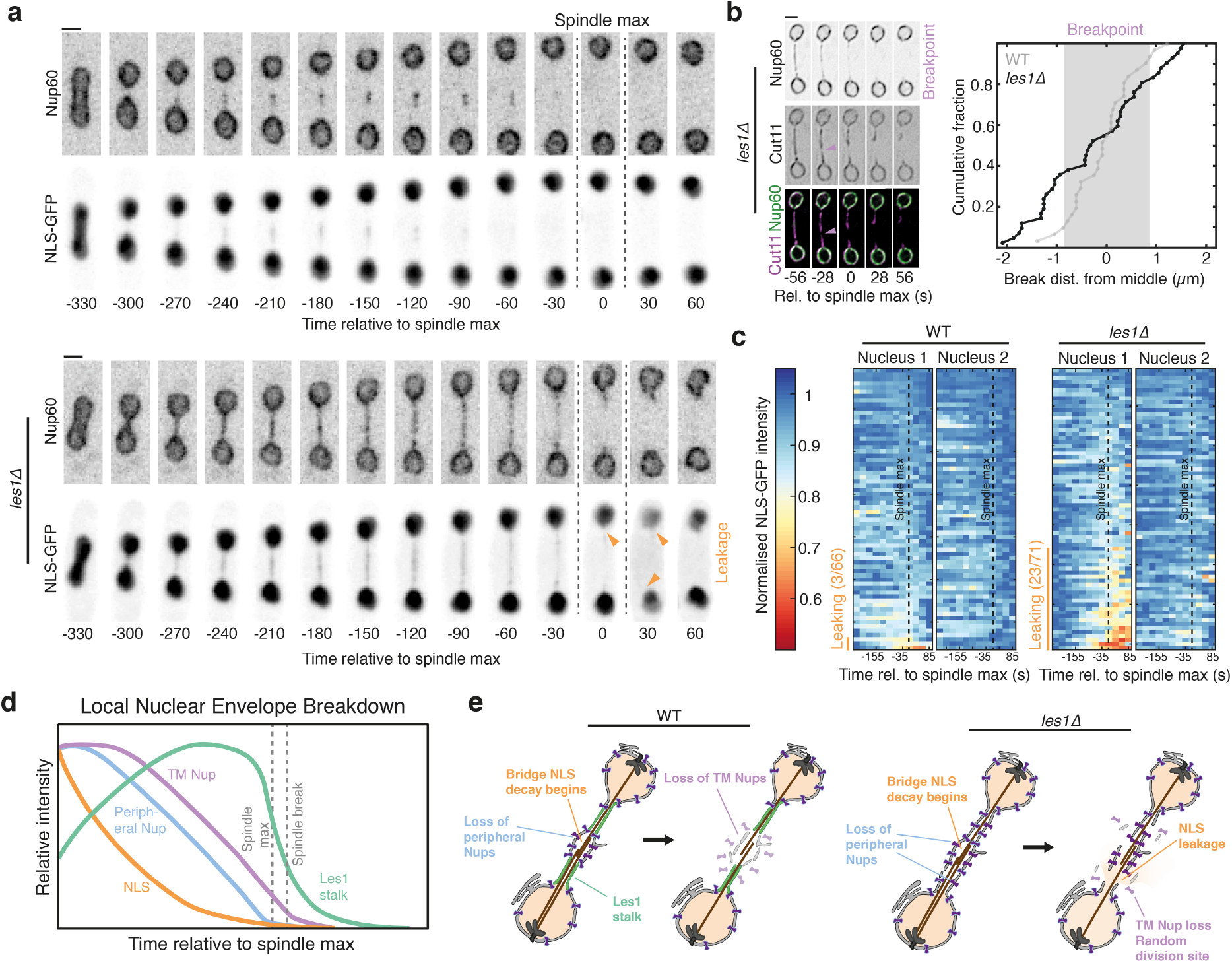
Les1 protects nuclear integrity during local NEB. a. Maximum intensity projections of spinning disk confocal images of cells expressing Nup60-mCherry tagged at the endogenous locus and NLS-GFP, either wildtype (top panels) or carrying a Les1 deletion (lower panels), imaged at 30 second intervals. Orange arrows indicate sites of NLS-GFP leakage from one or both daughter nuclei. Scale bars = 2 µm. b. SRRF reconstructions of confocal slices at 28 second intervals of *les1*Δ cells expressing Nup60-mNeonGreen and Cut11-mCherry tagged at the endogenous loci, aligned relative to spindle max (t=0). Magenta arrow indicates breakpoint. Scale bar = 2 µm. On the right, the cumulative distribution (42 cells from 2 strains) of breakpoint locations relative to the midzone in *les1*Δ cells. The shaded gray area represents the mean +/− standard deviation of breakpoint locations in wild-type cells, with the cumulative distribution as a gray line. c. Single cell intensity traces for wildtype (left 2 panels) and *les1*Δ cells (right 2 panels), indicating intensity in each daughter nucleus, aligned at spindle max (dotted line, t =0) and normalised individually to maximum intensity. Yellow bar and text indicate the number of leaky nuclei. d. Schematic indicating relative intensity levels of key readouts of the nuclear division process, aligned relative to spindle max. e. Schematic illustrating the process of local NEB and the role of Les1 in the spatial organisation of the bridge.

At the same time, daughter nuclei in the *les1*Δ strain were seen to suffer transient leakages of NLS-GFP close to the time of maximum spindle elongation (Figure 4a and 4c). These typically occurred in one of the two daughter cell nuclei (Figure 4a and 4c). Taken together, these data suggest that Les1 performs two roles when it accumulates in stalks in the nuclear bridge during mitotic exit. First, Les1 occludes NPCs to define a central region that is rich in NPCs, which is the site of later NPC disassembly, local NEB and thereby nuclear fission. Second, by helping to induce the formation of stalks, in which microtubules appear enclosed in a tight sheath of highly curved nuclear envelope, Les1 prevents the leakage of material from daughter cell nuclei. This ensures that while nuclei are topologically open to the cytoplasm at this stage of mitosis, the compartment boundary itself remains effectively closed.

## Conclusions

While we do not know precisely how Les1 fulfils these two roles, part of the explanation may be that Les1 generates a separated domain within the stalk membrane. This fits with the notion that in plants LEA proteins confer desiccation tolerance upon subcellular organelles (20) via the reorganization of intrinsically disordered domains. However, even then, the membrane at the tip of the two stalks must be sealed, where microtubule remnants project into the cytoplasm. We speculate that this may be achieved by the association of Les1 with ESCRTIII proteins (21), which have been implicated in NE sealing (22–24) and repair (25, 26) in both open and closed mitosis (in metazoans and fungi), because ESCRTIII genes were found to be the strongest negative genetic interactors of Les1/SPAC23C4.05c in a genome-wide genetic interaction screen (27).

Finally, we note that while nuclear fission occurs in Les1Δ cells at the wrong place, it does so at the right time (Figure S4a-b). Thus, there must be other cues, triggered by exit from mitosis, that lead to the gradual loss of NPCs from the bridge (but not from the two daughter nuclei), leading to local NEB and nuclear fission.

In summary, in this study we identify a protein Les1 that positions the site of nuclear fission during mitotic exit. At the same time, through the study of Les1 localisation and its deletion mutant, we reveal a close similarity between open and closed mitosis. In both cases, the new nuclear compartment is remodelled as the result of NPC disassembly. Thus, the key difference between mitotic strategies across the eukaryotic tree (28) may only be one of degree, depending on the timing and localisation of NPC disassembly.

## ACKNOWLEDGEMENTS

We would like to thank Mohan Balasubramanian, Snezhana Oliferenko, Silke Hauf, Jürg Bahler and Paul Nurse for sharing *S. pombe* strains and plasmids; the Oliferenko and Nurse labs for sharing expertise and S. pombe protocols; James O. Patterson for the kind gift of pFA6a-mNeonGreen plasmids; David Albrecht, Ishier Raote, Agathe Chaigne and members of the Baum lab for feedback on this manuscript. G.D. was funded by a European Union Marie Sklodowska-Curie Individual Fellowship (704281-CCDSA). Si.C. and R.H. were supported by the UK BBSRC (BB/R021805/1; BB/S507532/1), the UK Medical Research Council (MR/K015826/1), and the Wellcome Trust (203276/Z/16/Z). Sc.C was supported by the Francis Crick Institute which receives its core funding from Cancer Research UK (FC001121), the UK Medical Research Council (FC001121), and the Wellcome Trust (FC001121). W.K. was funded by the Medical Research Council (MC UP 1201/8). B.B. was supported by UCL’s Institute for the Physics of Living Systems, the MRC LMCB, by the Wellcome Trust (203276/Z/16/Z) and by Cancer Research UK (C1529/A28276).

## DATA AVAILABILITY STATEMENT

The data that support the findings of this study and all custom software written for this study will be made available upon reasonable request. Upon peer-reviewed publication, all strains generated for and used in this study will be made available upon reasonable request. All custom software will be deposited in a central repository prior to peer-reviewed publication.

## Methods

### A. *S. pombe* culture

*S. pombe* cells were cultured using standard methods (1, 2) on solid (YES-agar) and liquid (YES) rich growth media (Formedium), at a growth temperature of 32°C unless stated otherwise. All experiments were performed in exponential growth at 32°C with at least 48 hours of growth (>20 generations) in liquid YES before plating for live imaging. For live imaging, uncoated 35-mm dishes with polymer coverslips (No. 1.5 coverslip, 180 µm thick, Ibidi) were first coated with 1 mg/mL soybean lectin (in water, aliquots stored at −80°C, Sigma-Aldrich) for 15 minutes. After washing away the excess lectin with fresh YES, cells drawn from exponentially growing liquid cultures were allowed to settle for 30 minutes in a minimum volume of 500 µL of YES. The entire plating volume was replaced with 1 mL of fresh YES prewarmed to 32°C prior to transfer to the microscope. For cell culture in electron tomography experiments, see section on correlative fluorescence microscopy and electron tomography. For actin depolymerisation experiments, Latrunculin A (Sigma-Aldrich) was added to YES at 5 µM (10 mM stock in DMSO).

### B. Plasmids and *S. pombe* strain construction

The full genotypes of all strains used in this study are described in Table S1. Strains generated specifically for this study were constructed using standard methods (1, 2) for gene editing and crossing. Gene deletions and tagging were performed as previously described (3) for PCR-based gene targeting, using standard primers designed with the Bahler lab web-interface scripts (http://bahlerlab.info/resources/), pFA6a vector templates carrying Hygromycin (Hph) or Kanamycin (Kan) resistance cassettes, and transformation using the Lithium Acetate method (4). Antibiotic-resistant clones generated by this method were verified by PCR of the gene locus being targeted as well as fluorescence microscopy, if applicable. The exception to the standard workflow was for the pFA6a-mNeonGreen vectors used in this study, which carry a non-standard linker upstream of the mNeonGreen coding sequence. Instead of the standard 20-mer (CGGATCCCCGGGTTAATTAA) forward linker, these require a 21-mer forward linker (GATTCTGCTGGATCAGCTGGC). The reverse linker remains unchanged. One new pFA6a vector derivative was generated for this study, replacing the mCherry coding sequence in pFA6a-mCherry:Hph with the coding sequence for the photo-switchable fluorescent protein mEOS3.2 (5) (Addgene) by standard restriction-digestion cloning (using restriction enzymes BamHI and AscI, NEB). This vector is available upon request. Crosses were performed by random spore analysis (1, 2) followed by marker selection (Hygromycin/Kanamycin resistance, ura/leu auxotrophy, or sensitivity to 5 µM 1NM-PP1 (Calbiochem) for strains carrying the Ark1-as3 allele, as appropriate) followed by additional screening for fluorescence, if applicable.

### C. Live-cell fluorescence microscopy

All strains were imaged live in regular growth medium (YES) in glass-bottom dishes (see *S. pombe* culture) within stage-top incubation chambers held at 32°C. No single dish was used for experiments lasting longer than 3 hours from the time of plating. Three microscopes were used for this study: 2 spinning disk confocal systems and a Zeiss LSM880 with an Airyscan module. The first spinning disk microscope consists of a Nikon TiE inverted stand attached to a Yokogawa CSU-X1 spinning disc scan head and a Hamamatsu C9100-13 EMCCD camera. The entire system is controlled using Volocity software. Cells were imaged using a 100X oil-immersion CFI Plan Apochromat VC objective (1.4NA, working distance 0.13 mm) with an optional 1.5x tube lens. The second spinning disk microscope consists of a Zeiss AxioObserver Z1 inverted stand attached to a Yokogawa CSU-W1 spinning disc scan head and a Photometrics Prime BSI Scientific CMOS detector. Cells were imaged using 63X oil-immersion Plan Apochromat (1.4NA, working distance 0.19 mm) and 100X oil-immersion Plan Apochromat (1.4NA, working distance 0.17 mm) objectives combined with an optional 1.5x tube lens. The LSM880 is a combined confocal and multiphoton with an inverted Axio Observer microscope stand. Cells were imaged using a 63x oil-immersion Plan Apochromat objective (1.4NA, working distance 0.19 mm) and the Airyscan detector. Acquisition on the latter systems is controlled via the Zen software (Zeiss). In all cases, samples were illuminated with 488 nm (mNeonGreen or GFP) and 561 nm (mCherry) lasers and standard emission filter sets. Photoconversion of mEOS3.2 was carried out using a 405 nm laser. For regular live imaging, asynchronous cells were usually imaged using a 4.3 µm Z-stack with 16 slices at 270 nm vertical intervals, and time intervals ranging from 5 seconds to 120 seconds, never exceeding 30 minutes of continuous imaging. For Airyscan imaging, cells were imaged using a larger Z-stack at single timepoints. For SRRF and Hough fitting, cells were imaged with the system held at a single Z-plane at >3 frames per second.

### D. Image processing

All basic image processing (cropping, viewing stacks, scaling for visual presentation, producing maximum intensity projections) was carried out in Fiji (ImageJ) (6). All time-lapse images subjected to Super-Resolution Radial Fluctuations (SRRF) analysis were processed with NanoJ-LiveSRRF, the newest implementation of NanoJ-SRRF (7). NanoJ-LiveSRRF is available on request, expected to be available for download soon. NanoJ-SRRF is already released and available as open-source. Airyscan processing was carried out using proprietary Zen software (Zeiss).

### E. Analysis framework for single-cell trajectories

Analysis of single-cell trajectories was carried out using custom software written using the open-source platform Fiji (6).

#### E.1. ROI selection

Regions of Interest (ROIs) containing dividing cells were manually selected in time series data. Only division events that completed were selected for analysis (Figure S1b, ‘Manually select ROIs’, ‘Extracted ROI’). An ROI containing no nuclei was also selected for background subtraction of measured intensities.

#### E.2. Detecting divisions

Each ROI was maximum-intensity projected and this projection was then blurred, binarized, hole-filled and skeletonized using inbuilt Fiji (6) functions. The longest skeleton was assumed to correspond to the dividing nucleus, and the angle formed by this skeleton measured to be the division angle, *θ* (Figure S1b, ‘Determine division angle’).

#### E.3. Circle detection

Within ROIs, nuclei were identified for each frame and their radii determined using a custom-written Fiji plugin implementing the circular Hough transform (8) (Figure S1b, ‘Circle detection’). For two-colour images, the mNeon-Green channel was used to identify nuclei due to the superior signal-to-noise ratio, and these nuclei coordinates were assumed to be the same across both channels. For NLS images, a Sobel filter was used to highlight the nucleus perimeter prior to performing the circular Hough transform.

#### E.4. Identification of dividing nucleus pairs

For all possible pairs of detected circles in the ROI, the angle between the circle centres was calculated. Circle pairs at angles from *θ* by more than 30° were rejected. For proteins distributed along the whole length of the bridge between the two daughter nuclei (e.g. Les1 as shown in Figure 1d), candidate bridge paths were identified in each ROI frame by blurring, binarising, hole-filling and skeletonising the images (Figure S1b, ‘Path segmentation’). Paths of length < 3 pixels were rejected as these corresponded to isolated nuclei. The endpoints of the remaining paths were then checked against the coordinates of the remaining circle pairs. Paths without anchoring circle detections were rejected, as were any detected circles lacking an associated path. The final result for each frame was either that no divisions were detected, or two nuclei and joining path were detected (Figure S1b, ‘Filter circles and paths satisfying selection criteria’). Following breakage of the bridge, there is no longer a complete path to between the daughter nuclei. In this case, paths with one endpoint anchored in two detected circles at an appropriate angle were joined by a straight line between the ‘free’ endpoints of the two paths (Figure S1b, ‘Filter circles and paths satisfying selection criteria’, yellow dotted line). For cases where the protein was not present along the whole length of the bridge (e.g. NLS as shown in Figure 1a) the path between the two nuclei was defined as a straight line between the two circle centres.

#### E.5. Definition of bridge and timepoints

The bridge was defined as the path between the two daughter nuclei excluding any pixels within the nuclei perimeters. The initiation of division was defined as the first frame in which two nuclei were successfully detected, and dumbbell appearance was defined as the first frame where a bridge could be discerned (i.e. the first timepoint where all path pixels were not contained by daughter nuclei). Nucleus separation was defined as the Euclidean distance between the centroids of the daughter nuclei.

#### E.6. Measurement of nuclear intensities

Nuclear intensities were the background-subtracted average intensities within the detected circles (Figure 1b ‘Nucleus’, Figure 4c).

#### E.7. Measurement of bridge intensities

Total bridge intensity was the background-subtracted average intensity along the bridge path (Figure 1b ‘Bridge’, Figure 3g, Figure S4). Stalk intensities were the background-subtracted average intensities for bridge path pixels within 1.5 µm of a nucleus perimeter (Figure 1e, Figure S1e ‘Stalk’). Midzone intensities were the background-subtracted average intensities for bridge path pixels more than 1.5 µm away from a nucleus perimeter (Figure S1e, ‘Midzone’). In all cases, the path linewidth was set to 5 (i.e. 2 pixels perpendicularly either side of the path) to account for the full thickness of the bridges.

#### E.8. Measurement of vertical displacement of NE proteins

Circle detection and analysis was again performed using the circular Hough transform, but this time on whole frames (i.e. no manually selected ROIs) so that all nuclei within a single time frame were detected. All data analysed had Cut11 as a reference strain in one channel and another NE protein of interest in the second channel. For each nucleus, the difference in radius between the two channels was calculated. SRRF processing was performed on images prior to radius measurement to increase resolution. As a control, Nup60 was labelled with a green-to-red photoconvertable fluorescent protein (mEOS3.2) and images taken before and after photoconversion. These two channels were then analysed using the same pipeline as for the Cut11 two-colour strains to check that there were no systematic errors in radius measurement between red and green channels. The values obtained for various Nups provide a good match with a recent electron microscopy analysis of the *S. pombe* NPC (9), which also provided the estimate of the width of the lumen between inner and outer nuclear envelopes.

#### E.9. Spindle detection and breakpoint measurement

Spindles were segmented from ROIs in images containing fluorescently-labelled tubulin (mCherry-Atb2) by blurring, binarisation, hole-filling and skeletonisation. The longest skeleton was determined for each frame; the frame in which spindle breakage occurred was determined as the first frame where the maximum skeleton length decreased by *≥* 25% compared to the maximum skeleton length measured across all previous frames.

#### E.10. Data curation

Every detected nucleus division was manually checked to ensure that the correct nuclei had been identified, and ROIs containing false detections were excluded from analysis.

#### E.11. Manual quantification

The breakpoint analysis in Figure 4b was scored manually-due to the high error rates of identifying the precise site of breakage from automatically extracted time series data. The timings in Figure S3d-f (with reference to the first appearance of the nuclear bridge) were manually quantified. This was due to the presence of a second green label (Cdc15-mNeonGreen) in addition to Cut11-GFP: while this allowed the visualisation of the cytokinetic ring, it also prevented the automated analysis of maximum intensity projections.

### F. Population-level analyses and statistics

Unless otherwise specified, single-cell trajectories (see section above for specific measurements) were aligned to the time of maximal spindle elongation (if a tubulin label not present, measured indirectly using the maximal separation between daughter nuclei as a proxy for spindle length) – making use of the fact that spindle elongation kinetics are highly reproducible from cell to cell. When combining trajectories from different strains (e.g. Figure 3g), we made use of the observation that mitotic timing tends to be consistent even when spindles reach different maximum lengths (Figure S3b). This is probably due to a Klp9-dependent adjustment in spindle elongation rates between strains (strains with longer spindles elongate their spindles faster) (10). Statistical analyses were carried out using GraphPad (mean and standard deviation of averaged traces), MATLAB (ANOVA; mean and standard deviation of averaged traces) and the Estimation Stats platform (11) (two-sided Mann-Whitney). Graphs and heatmaps were generated using either GraphPad or MATLAB.

### G. Correlative fluorescence microscopy and electron tomography

Correlative microscopy was done as described before for resin-embedded yeast cells (12, 13), with minor modifications. In brief, Les1-mNeonGreen/mCherry-Atb2 expressing cells were grown in YES at 32°C to mid-log phase, pelleted by vacuum-filtration and high-pressure frozen in the 100 µm recess of aluminium platelets (Wohlwend) using an HPM100 (Leica Microsystems). Samples were freeze-substituted and embedded in Lowicryl HM20 (Polysciences) according to the published protocol (12), except with 0.03 uranyl acetate in the freeze-substitution solution. Resin blocks were sectioned at 320 nm nominal thickness, picked up onto carbon-coated copper grids (AGS160, Agar Scientific) and imaged on the same day on a Nikon TE2000 microscope using a 100x TIRF objective, a NEO sCMOS DC-152Q-C00-FI camera (Andor), a Niji LED light box and filter sets 49002 ET-EGFP (Chroma) for mNeonGreen signals and 49005 ET-DSRed (Chroma) for mCherry. Based on the fluorescence images, cells profiles in which an elongated bridge was visible within the section plane were selected for electron tomography. 15 nm protein A-coated gold beads (EMS) were adhered to the grids prior to Reynolds’ lead citrate staining. Dual-axis electron tomographic tilt series were acquired approximately from +60° to −60° on a TF20 electron microscope (FEI) operated in STEM mode, using a 50 µm C2 aperture, at 1° increment and 1.1 nm pixel size on an axial bright field detector (14), using SerialEM (15). Tomograms were reconstructed using IMOD (16). Segmentation models were generated using Amira (Thermo Fisher Scientific) by manual tracing of membranes and microtubules. Electron tomographic slices shown in figures have been mildly gauss-filtered to improve visibility.

**Table S1.**
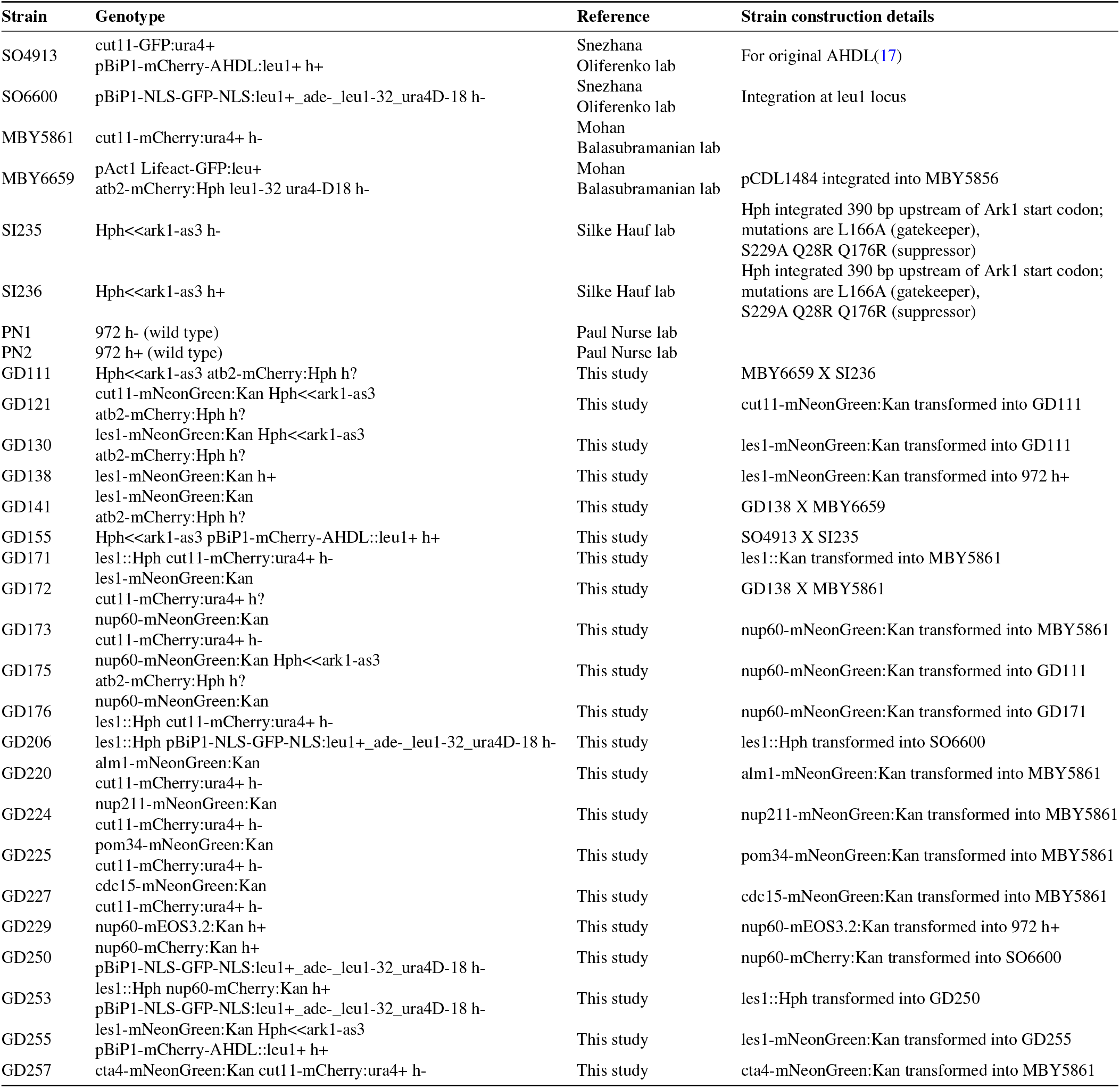
Complete list of *S. pombe* strains used in this study.

---

**Fig. S1.**
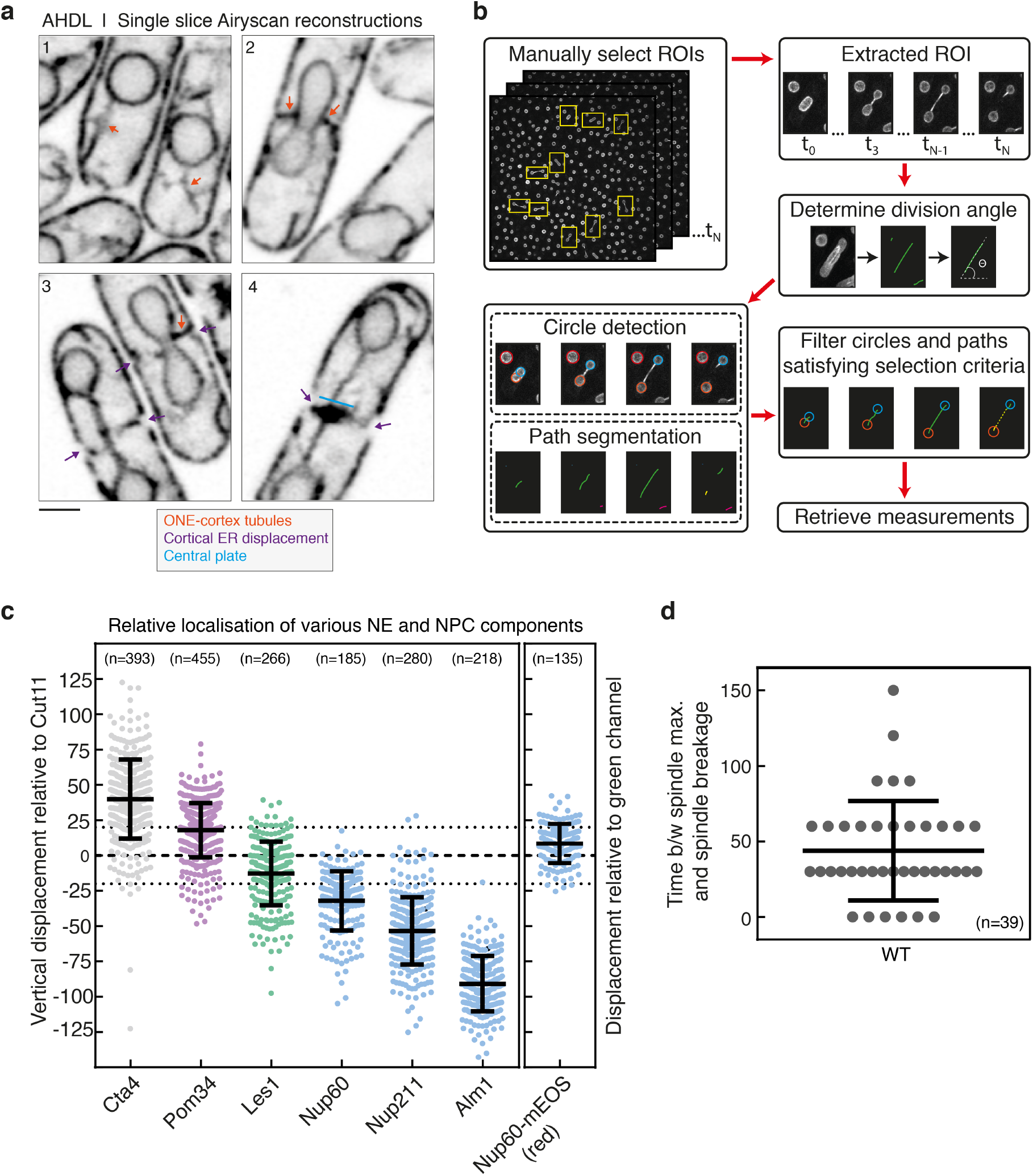
a. Airyscan reconstructions of cells expressing the mCherry-AHDL synthetic construct. Orange arrows indicate tubules linking the outer nuclear envelope to the cortex. Purple arrows indicate the displacement of the cortical ER by the division ring (not shown). The blue bar indicates the central ER plate formed during late anaphase. Scale bar = 2 µm. b. Schematic demonstrating pipeline for detecting and measuring nuclei and bridges in timelapse data. Individual steps are described fully in the Methods. Representative images shown here are of Les1-mNeonGreen and Atb2-mCherry. c. The vertical displacement (relative to the plane of the nuclear envelope) of various nuclear pore complex (Nup60, in the nucleoplasmic ring; Alm1 and Nup211 in the basket; transmembrane Nup Pom34) and NE membrane proteins (Cta4) relative to Cut11. These measurements were made by fitting a circular Hough transform to SRRF-processed 2-colour images of interphase nuclei (see Methods for details). Nup60-mEOS was used as an internal control, with the values representing the displacement of photo-converted Nup60-mEOS (red channel) relative to un-converted Nup60-mEOS (green channel). The dotted lines represent an estimate of the thickness of the nuclear envelope (see Methods for details). d. The delay between reaching maximum spindle length (spindle max.) and spindle breakage, in seconds. Spindle breakage always follows maximal extension.

**Fig. S2.**
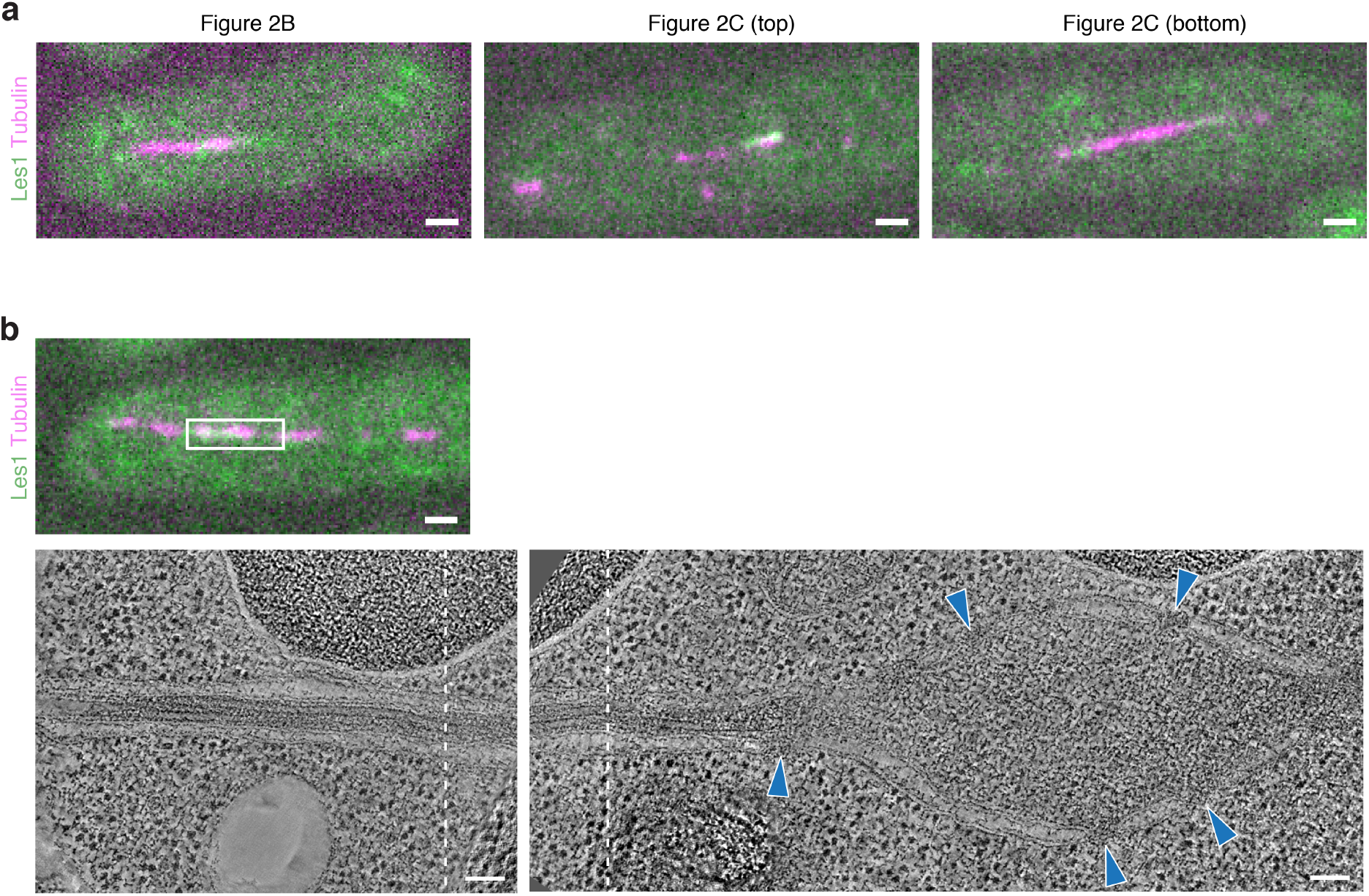
a. Fluorescence images of resin section through cells expressing Les1-mNeonGreen (green) and Atb2-mCherry (magenta), corresponding to cells that were imaged by electron tomography and are shown in Figure 2b (left image), Figure 2c top panel (middle image) and Figure 2c lower panel (right image). Images have been rotated to match approximately the orientation of electron tomograms. Scale bars = 1 µm. b. Fluorescence image of resin section through cell expressing Les1-mNeonGreen (green) and Atb2-mCherry (magenta), corresponding to cell imaged by electron tomography. White region corresponds to regions shown in lower panel. Scale bar = 1 µm. Lower panel are virtual slices through electron tomograms of the cell shown above. The approximate overlap in field of view of the tomograms is indicated by dashed lines. Note that no NPCs are visible in the stalk part shown in the left image. NPCs (indicated by blue arrowheads) are constrained to the midzone (right image). Scale bar = 100 nm.

**Fig. S3.**
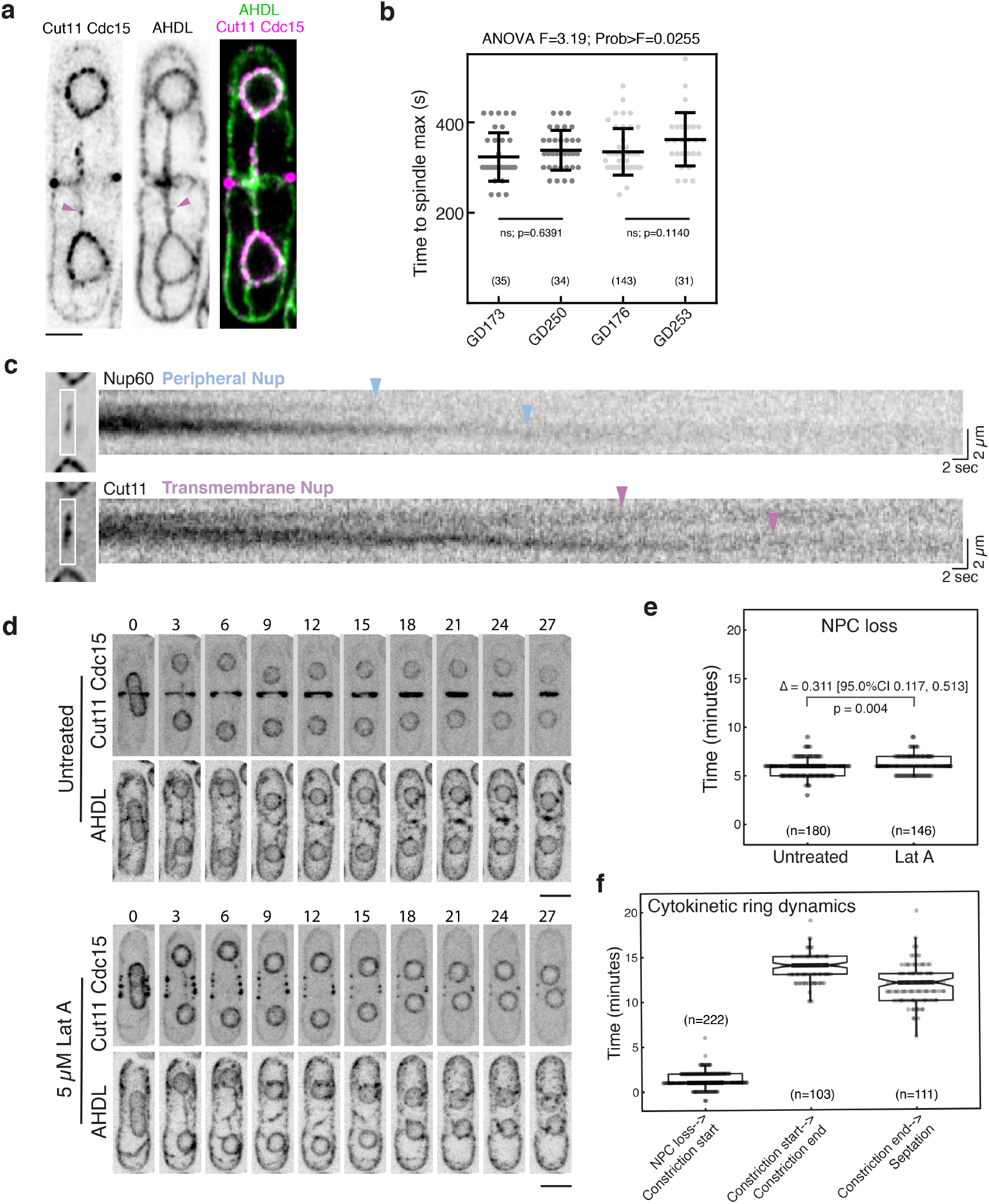
a. Single Airyscan reconstructions of cells expressing Cut11 tagged at the endogenous locus with mNeonGreen and a synthetic mCherry-AHDL construct. Magenta arrow highlights stray nuclear pore cluster accompanied by widening of the nuclear envelope. Scale bar = 2 µm. b. Time from bridge formation to maximum spindle length (c) measured for strains pooled to generate Figure 3g (GD173, GD250) and S4a (GD176, GD253). Numbers in brackets indicate number of cells in each population, with bars representing mean and standard deviation. The ANOVA F statistic and p-value are listed above each plot. The line and pairwise p-value within each plot refer to the comparison between pooled strains. c. Kymograph generated using 10fps single plane imaging of a strain expressing Nup60 and Cut11 tagged at the endogenous loci with mNeonGreen and mCherry respectively. Blue (Nup60) and magenta (Cut11) arrows represent the staggered decay of individual clusters of nuclear pores. d. Timelapse confocal images of cells expressing Cut11-GFP, Cdc15-mNeonGreen, and mCherry-AHDL acquired at 60 second intervals with frames displayed at 3 minute intervals. Scale bars =4 µm. Treatment with Latrunculin A depolymerises the actin ring (marked by Cdc15) but has a minimal impact on the time of nuclear division, as marked by the time from bridge formation to complete NPC signal loss (e). Numbers above and below the horizontal bar represent the difference in means with 95% confidence interval and the two-sided Mann-Whitney p-value. Numbers in brackets represent the number of cells in each population. f. Cytokinetic ring constriction only begins after nuclear division completes, and it takes almost 30 minutes for the ring to constrict and septation to complete.

**Fig. S4.**
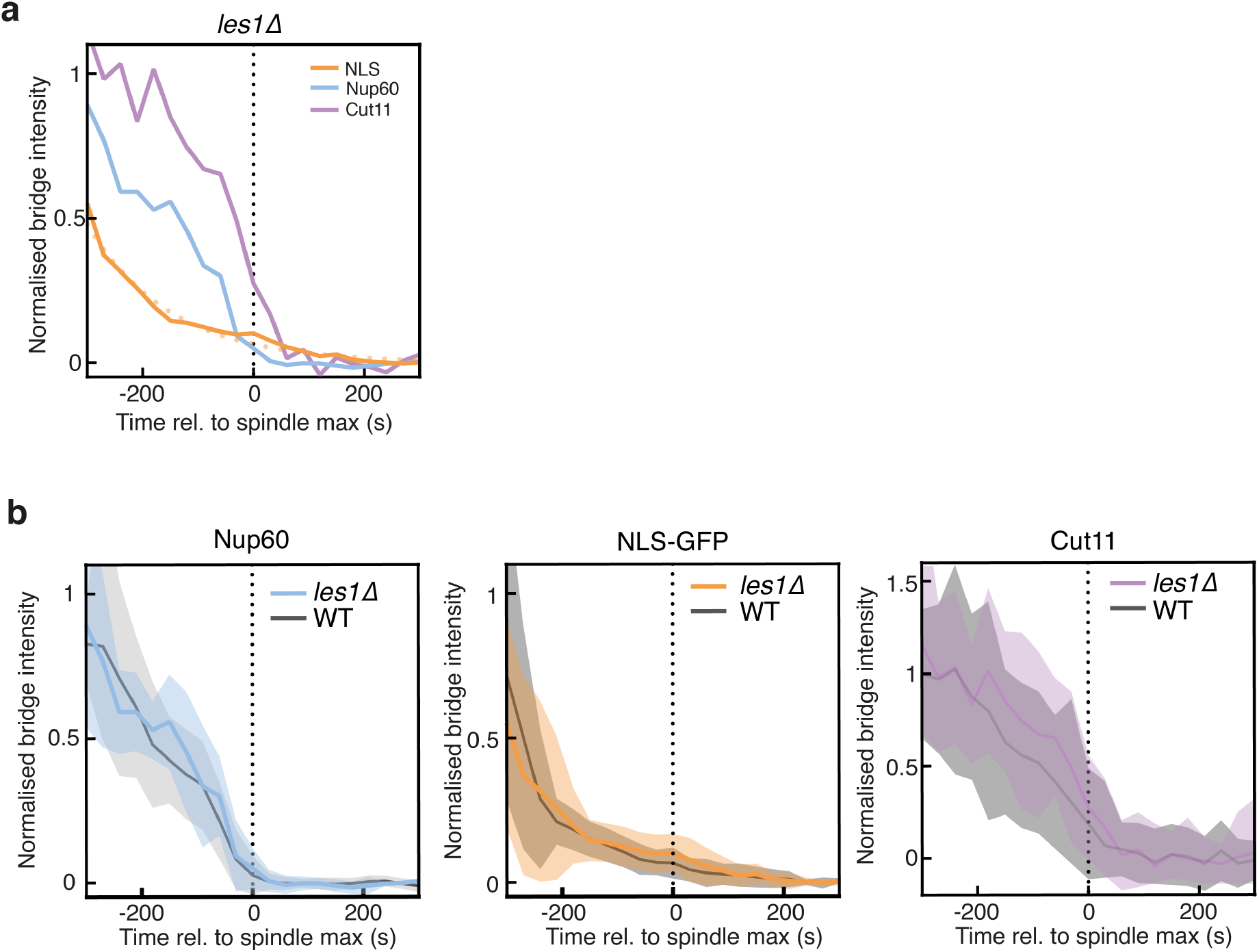
a. Mean decay curves for NLS-GFP (yellow, from 28 cells at t=-300 to 39 cells at t=0), Nup60 (blue, from 10 cells at t=-300 to 15 cells at t=0) and Cut11 (magenta, from 10 cells at t=-300 to 15 cells at t=0) in a les1Δ background. Each trace normalized by division by maximum bridge signal for that cell prior to averaging. Dotted yellow line indicates exponential fit to NLS-GFP average. b. Mean plus a band representing standard deviation for each marker in a. relative to wild-type cells (mean data for wildtype cells in Figure 3g).

